## Supplementary Figures for "Separating the control of moving and holding in post-stroke arm paresis"

### Supplementary Materials

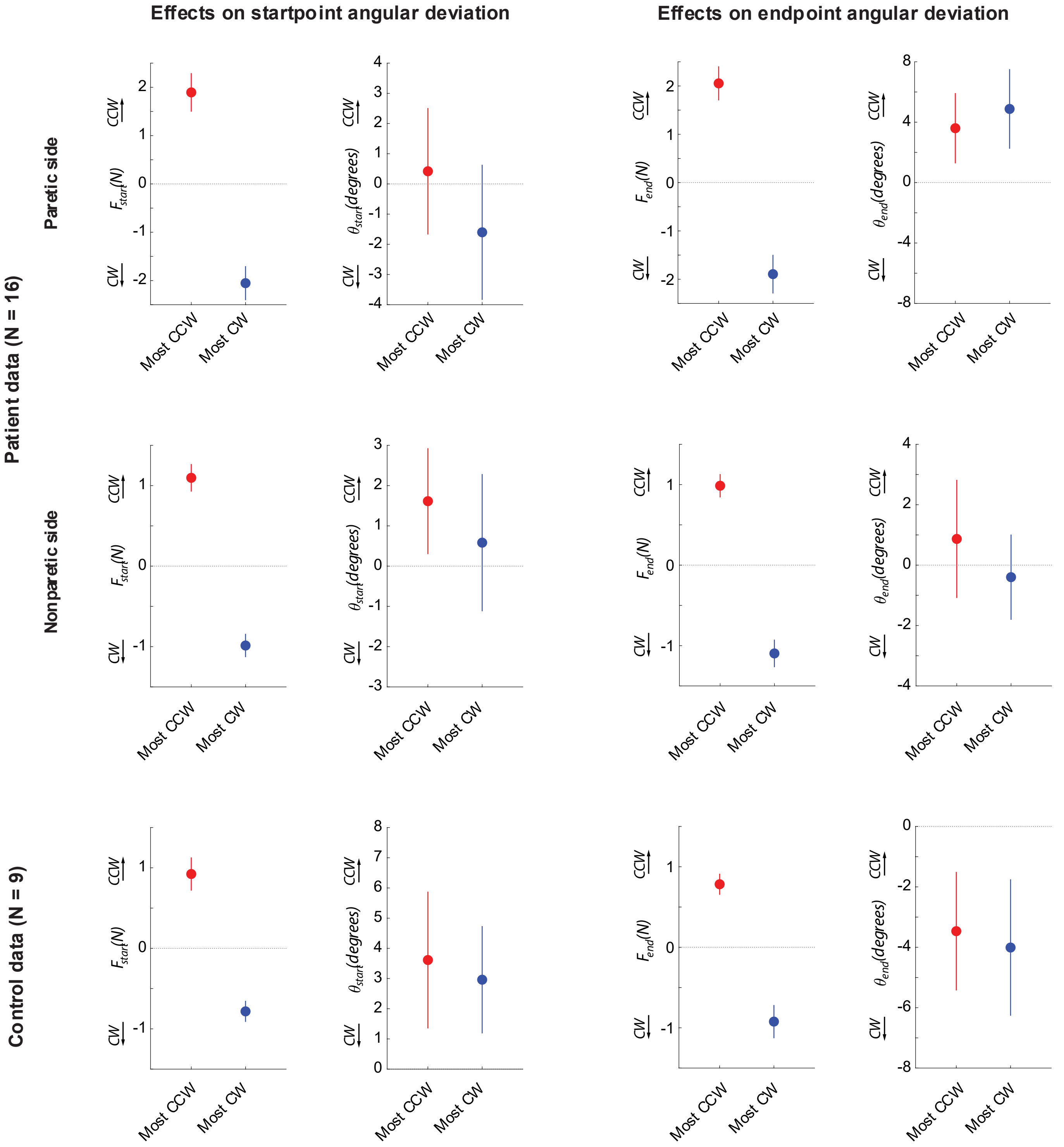


***Figure S5-1: Relationships between active reaching and resting posture biases for the non-paretic side of patients and for controls. A:*** *Reproduction of Figure 5E(right) and Figure 5F(right) for reference (data from the paretic side of stroke patients). Left: resting biases corresponding to the instances (target directions) where the component of the resting bias lateral to the target direction was the most CW (blue) vs. the most CCW-oriented (red). Right: corresponding initial angular deviations.* ***B:*** *Same analyses for the non-paretic arm of stroke patients.* ***C:*** *Same analyses for the control data.*

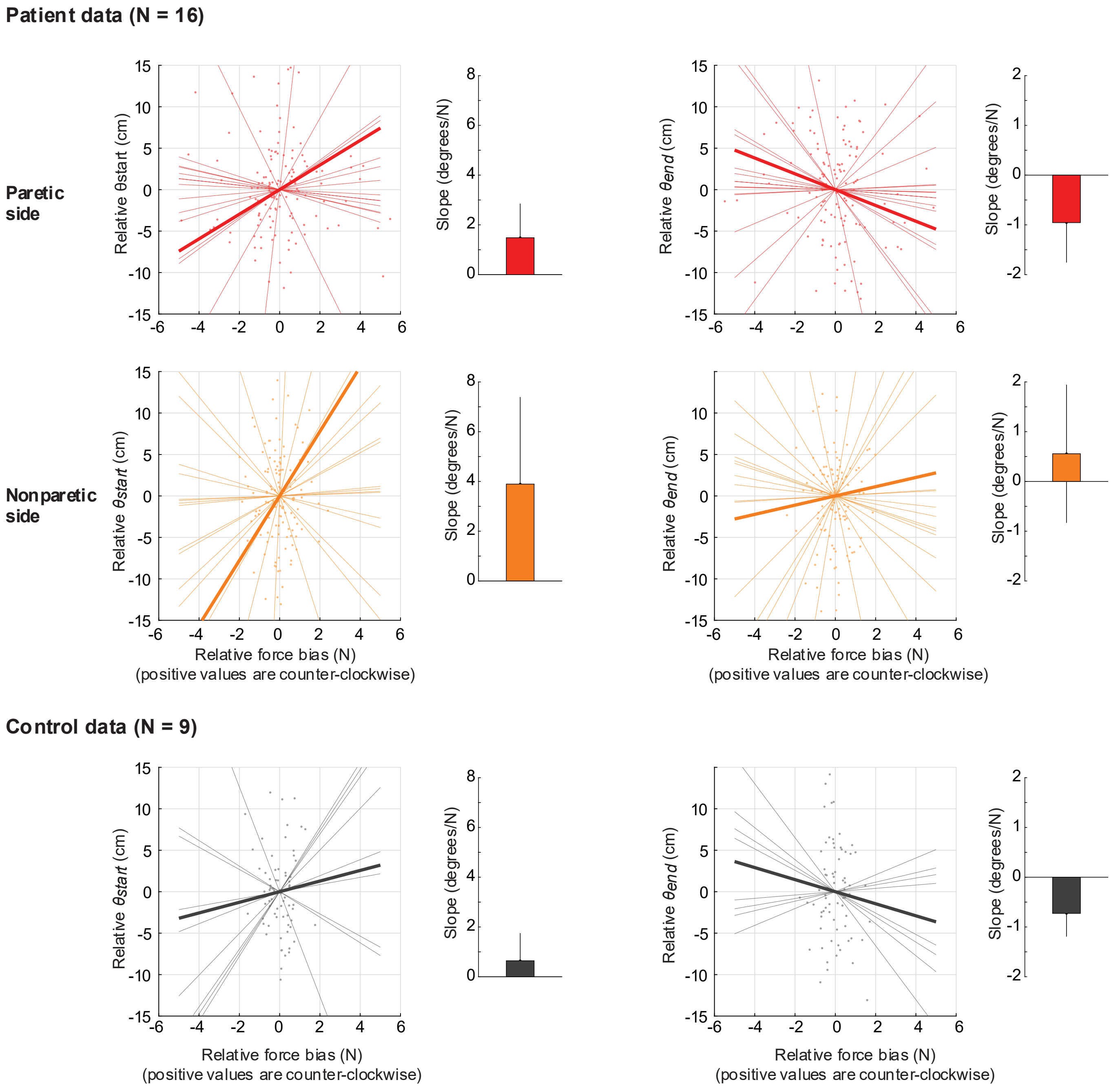


***Figure S5-2: Estimating the sensitivity of active reaching to resting posture biases.*** *This analysis, in contrast to the main analysis in Figure 5E/F which only compares the instances where resting biases are the most CW/CCW related to movement direction, takes all possible instances and uses linear regression to estimate the sensitivity (slope) of initial and endpoint angular deviations to resting biases on the start and target positions, respectively.* ***(A)*** *Analysis for the patients' paretic arm. Left: Shown is a scatter plot of the relative (i.e. mean-subtracted) initial angular deviation (θstart, y-axis) against the relative force bias lateral to movement (Fstart, x-axis). Each dot represents one movement direction for one participant. Both x- and y-axis data were mean-subtracted separately for each participant. The thin lines indicate linear fits for each participant; the thick line indicates the average of those fits. The bar graph to the right shows the average slope (sensitivity) ± SEM. Positive slopes indicate that angular deviations follow the posture bias at start; there is no evidence of a positive slope as shown in the bar graph, in line with our main analysis in Figure 5F. Right: same but for endpoint angular deviations (θend) against resting biases at the target (Fend).* ***(B) and (C)****: Same as in (A) but for the non-paretic arm of patients and for healthy controls, respectively.*

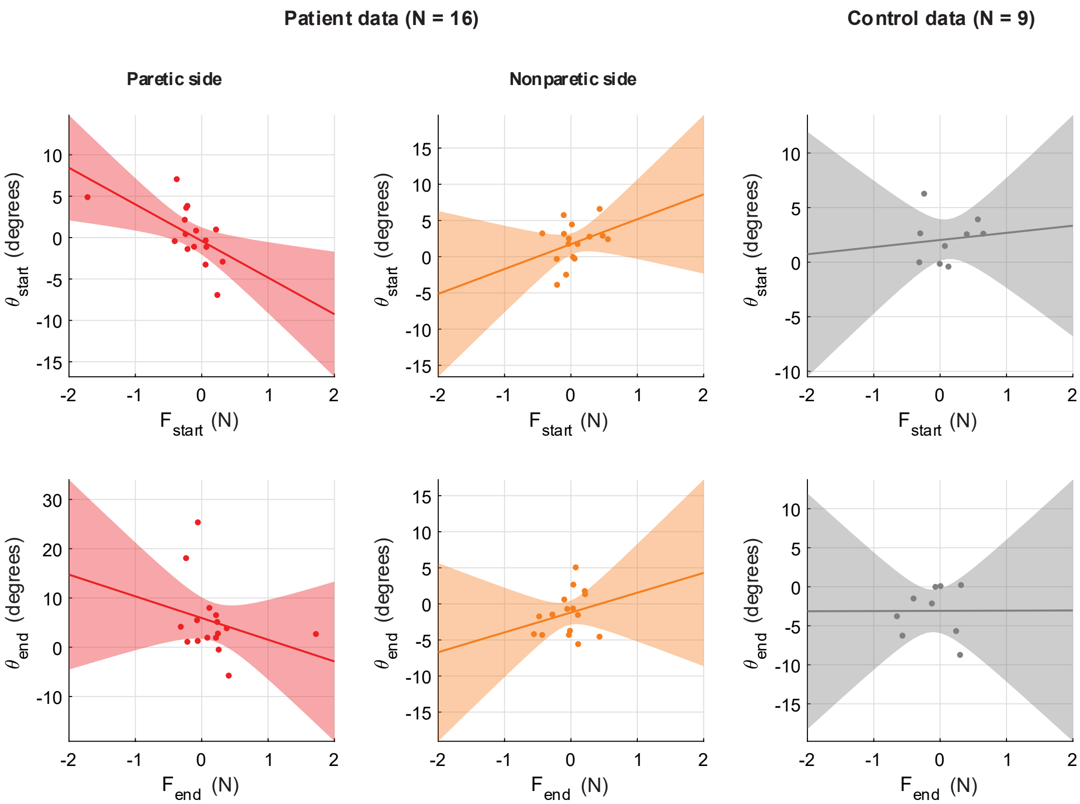


***Figure S5-3: Across-individual correlations between resting bias and movement kinematics. (A)*** *Scatter plot of average θstart the average component of Fstart lateral to target direction. Positive numbers indicate counter-clockwise θstart and Fstart. Solid line indicates linear fit; shading illustrates the associated 95% confidence interval. Plots indicate a lack of positive relationship. While this is in line with the main analysis, these across-individual relationships can only provide limited evidence since they may average out opposing contributions of resting biases. This is why our main analysis focuses on within-individual differences across different movement directions.* ***(B, C)****: Same as (A) but for the non-paretic side of patients* ***(B)*** *and for healthy controls* ***(C)****.* ***(D-F):*** *same as (A-C) but for approach angle close to the endpoint, θend.*

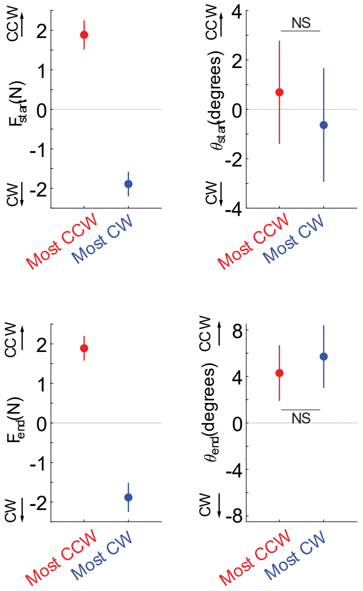


***Figure S5-E:*** *Repetition of the analysis in Figure 5E/F (top) and 5H/I (bottom) but with resting biases calculated without trial rejection, showing similar results (lack of relationship between θstart and θend with Fstart and Fend, respectively).*

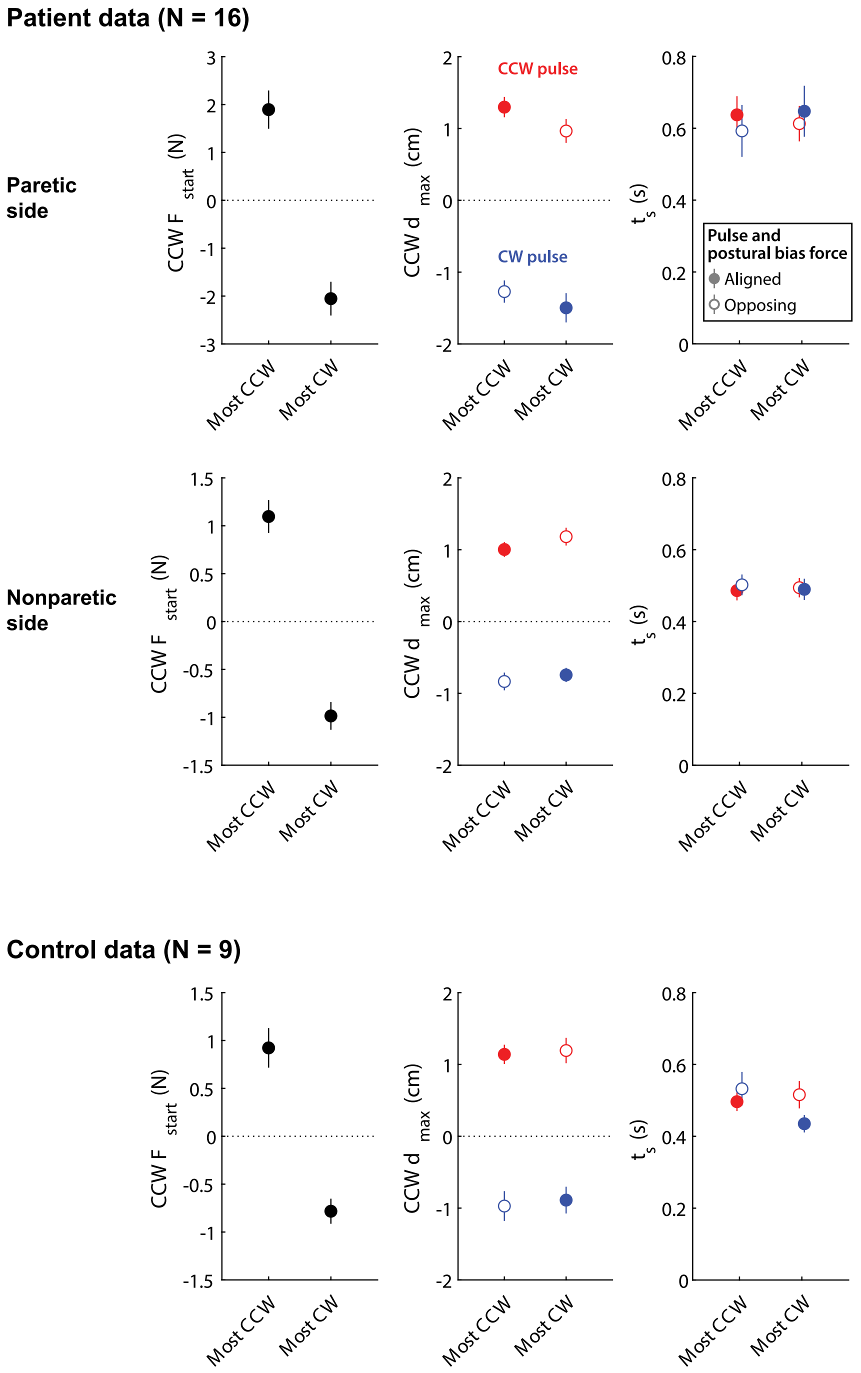


***Figure S6-1: Relationships between responses to pulse perturbations and resting posture biases for the non-paretic side of patients and for controls. Top row:*** *Reproduction of Figure 6D for reference (data from the paretic side of stroke patients).* ***Middle row:*** *Same analyses for the non-paretic arm of stroke patients.* ***Bottom row:*** *Same analyses for the control data. Note, in all cases, the lack of difference between the instances where postural biases were the most aligned vs. the most opposed to the pulse.*

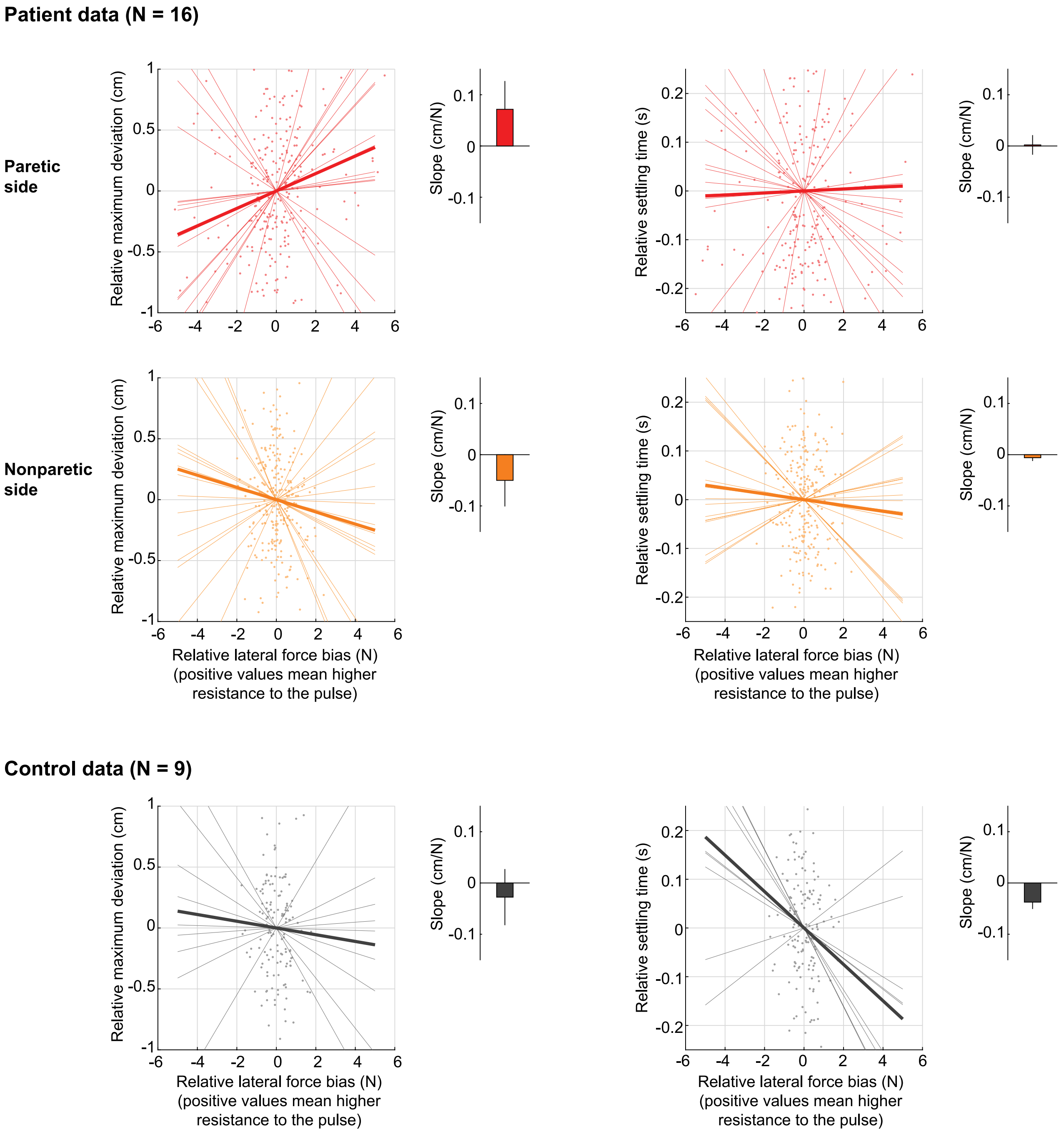


***Figure S6-2: Estimating the sensitivity of movement perturbation responses to resting posture biases.*** *This analysis, in contrast to the main analysis in Figure 6D which only compares the instances where resting biases are the most aligned vs. the most opposed to the pulse perturbations, takes all possible instances and uses linear regression to estimate the sensitivity (slope) of maximum deviation to resting biases.* ***(Top row)*** *Analysis for the patients' paretic arm. Left: Shown is a scatter plot of the relative (i.e. mean-subtracted) maximum deviation (y-axis) against the relative force bias in the perturbation direction (x-axis), with positive values indicating increased resistance to the pulse. Each dot indicates one movement direction/pulse sign combination for one participant. Both x- and y-axis data were mean-subtracted separately for each participant and pulse type. The thin lines indicate linear fits for each participant; the thick line indicates the average of those fits. The bar graph shows the average slope (sensitivity) ± SEM. Negative slopes indicate interact with responses to perturbations during active movement). There is no evidence of a negative slope as shown in the bar graph, in line with our main analysis in Figure 6D. Right: Same as left, but for the relative settling time.* ***(Middle row) and (Bottom row)****: Same as in (A) but for the non-paretic arm of patients and for healthy controls, respectively.*

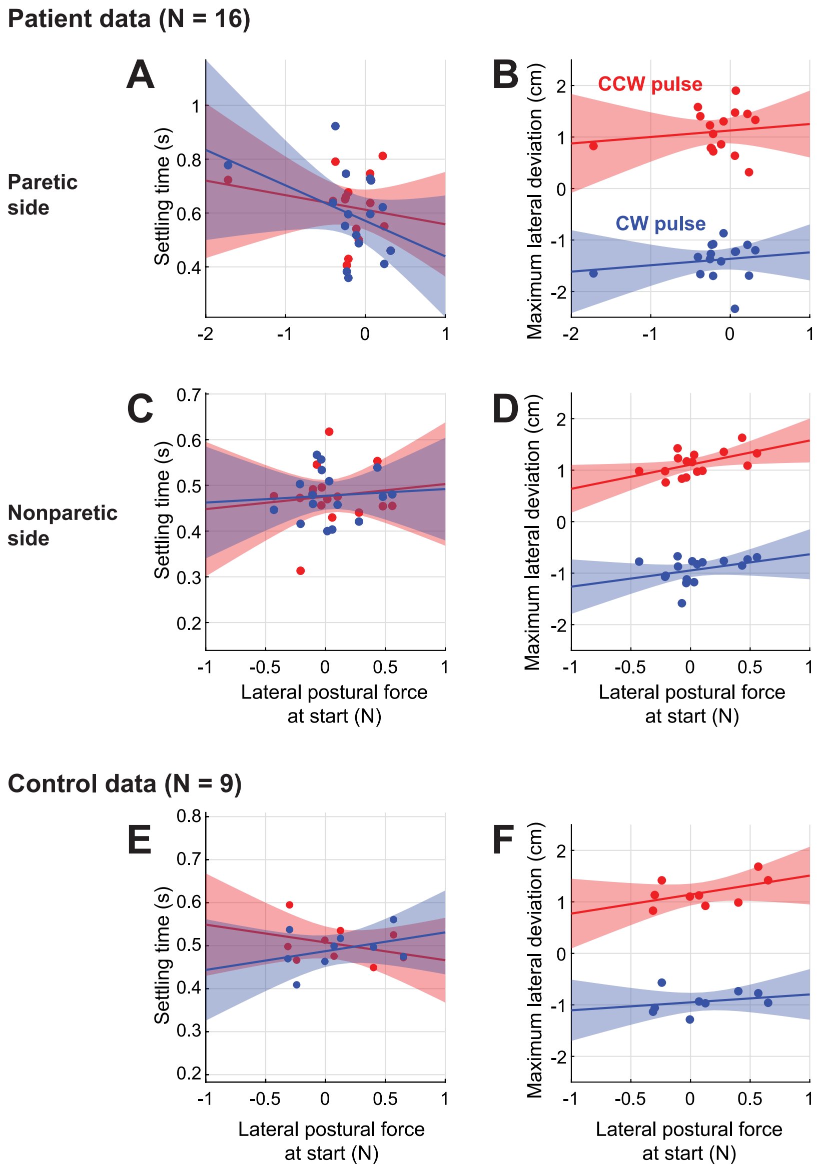


***Figure S6-3: (A)*** *Across-patient comparison between settling time and lateral postural bias forces on movement start. Paretic data shown. Red: CCW pulse; Blue: CW pulse.* ***(B)*** *same as (A), but for (signed) maximum lateral deviation for the two pulse types.* ***(C,D):*** *same as (A,B) but for nonparetic data.* ***(E,F):*** *same as (A,B) but for control data.*

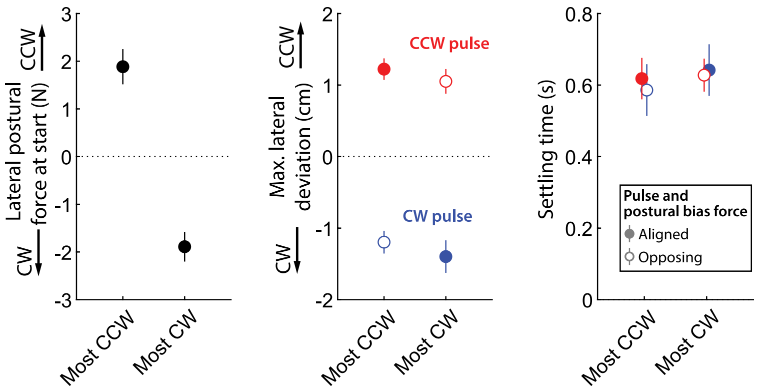


***Figure S6-E:*** *Repetition of the analysis in Figure 6D but with resting biases calculated without trial rejection, showing similar results (lack of effect of resting biases upon the response to the force pulse).*

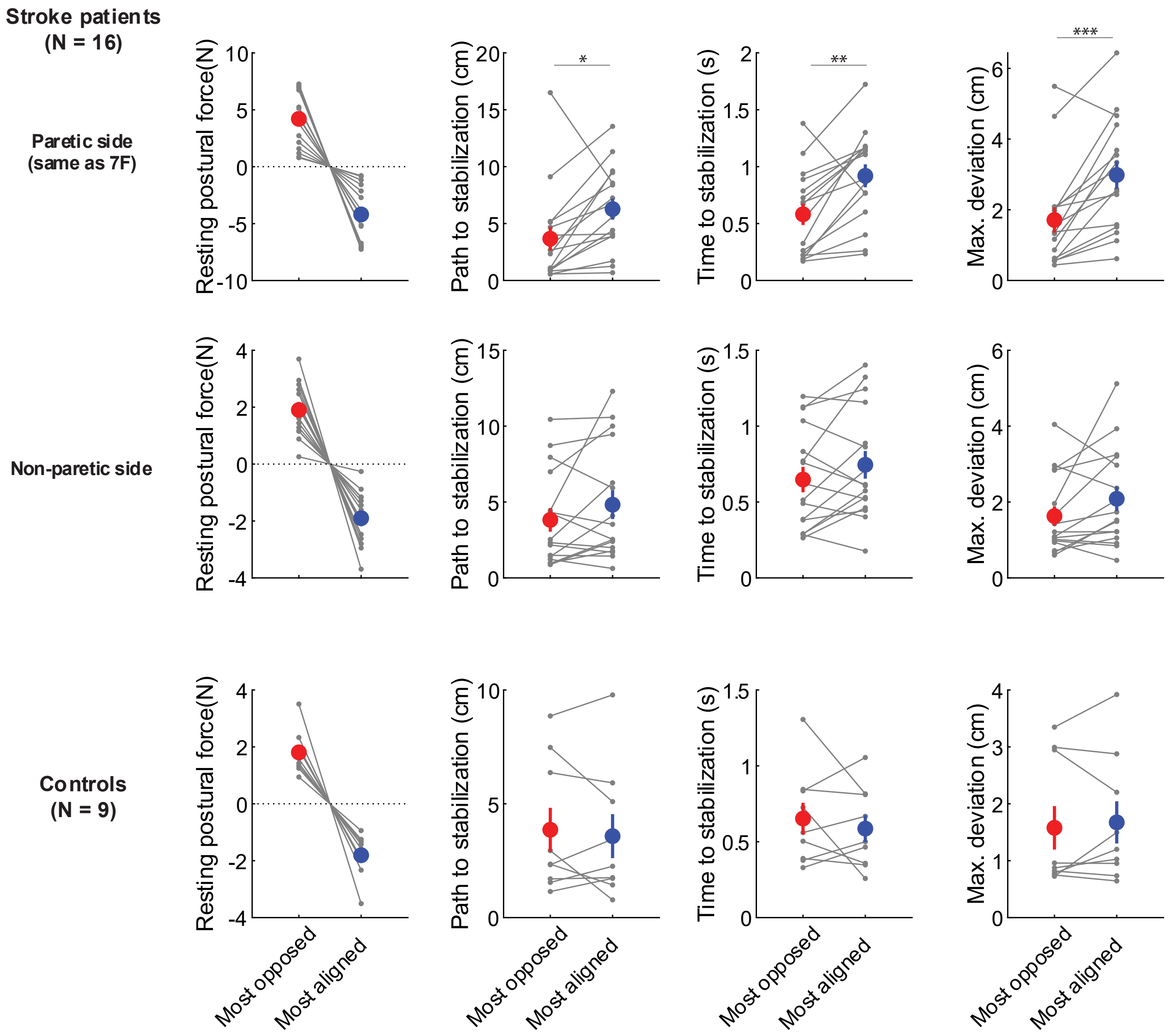


***Figure S7-1: Relationships between responses to static perturbations and resting posture biases for the non-paretic side of patients and for controls. Top row:*** *Reproduction of Figure 7F for reference (data from the paretic side of stroke patients).* ***Middle row:*** *Same analyses for the non-paretic arm of stroke patients. Note that, while the magnitude of the corresponding resting biases is lower compared to the paretic arm, the outcome variables (path to stabilization, time to stabilization, and maximum deviation) are nominally (but not significantly) lower in the case where resting biases are most opposed to the perturbation.* ***Bottom row:*** *Same analyses for the control data.*

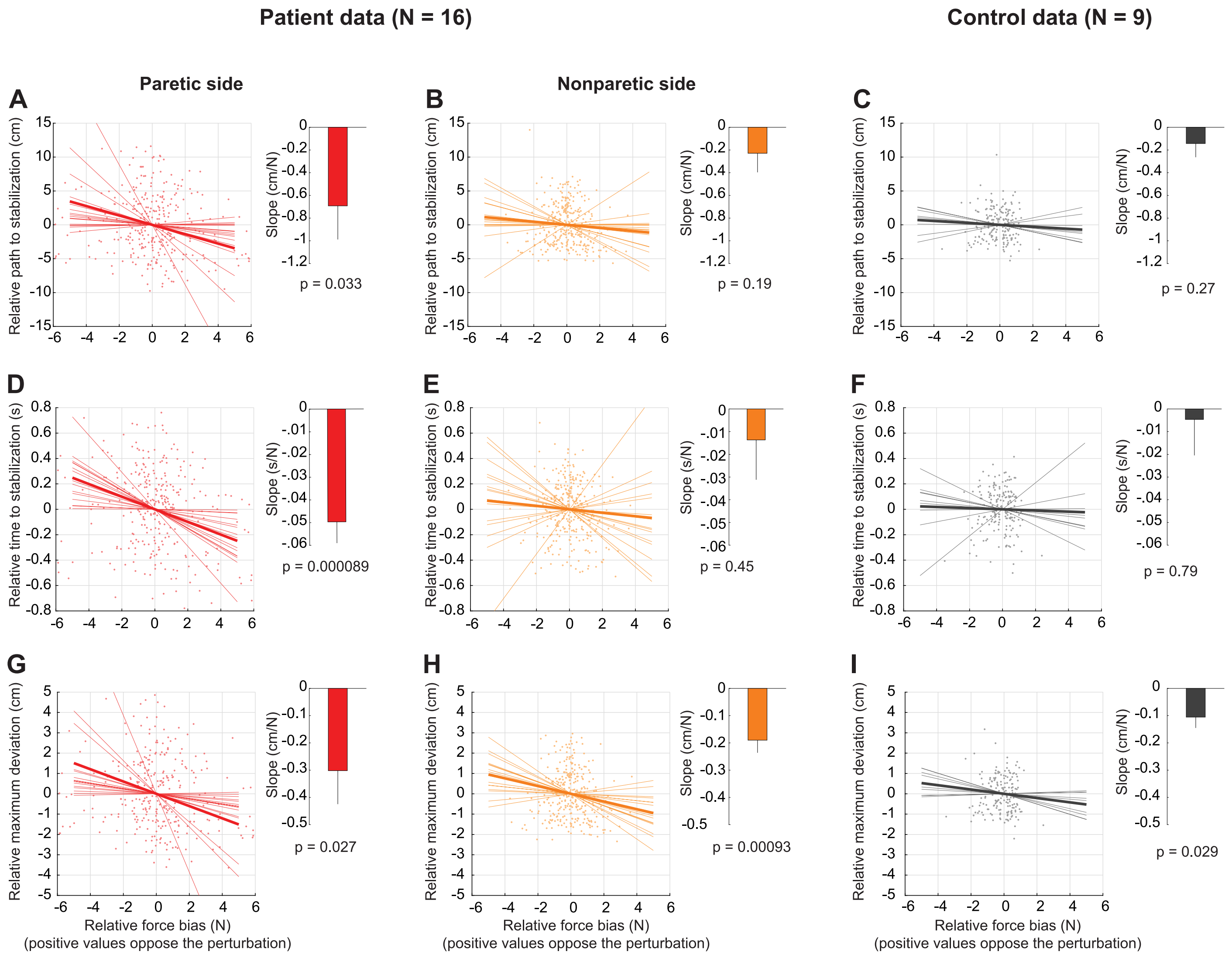


***Figure S7-2: Estimating the sensitivity of active holding control to resting posture biases.*** *This analysis, in contrast to the main analysis in Figure 7F which only compares the instances where resting biases are the most aligned vs. the most opposed to the static perturbation, takes all possible instances and uses linear regression to estimate the sensitivity (slope) of the outcome variables to resting biases.* ***(A)*** *Analysis for the patients' paretic arm. Shown is a scatter plot of the relative (i.e. mean-subtracted) path to stabilization (y-axis) against the relative force bias in the perturbation direction (x-axis). Both x- and y-axis data were mean-subtracted separately for each participant. The thin lines indicate linear fits for each participant; the thick line indicates the average of those fits. The bar graph to the right shows the average slope (sensitivity) ± SEM, which was, in this case, significantly negative in line with an effect of resting bias on patients' performance against the holding perturbation.* ***(B) and (C)****: Same as in (A) but for the non-paretic arm of patients and for healthy controls, respectively.* ***(D-F)****: Same as A-C but for relative time to stabilization.* ***(G-I)*** *Same as A-C but for relative maximum deviation. Note how all three metrics demonstrate that patients are more able to resist and recover from the static perturbation when resting biases are directed against the perturbation (A,D,G). In turn, this supports the evidence shown in Fig. 7F that resting biases interact with active holding control.*

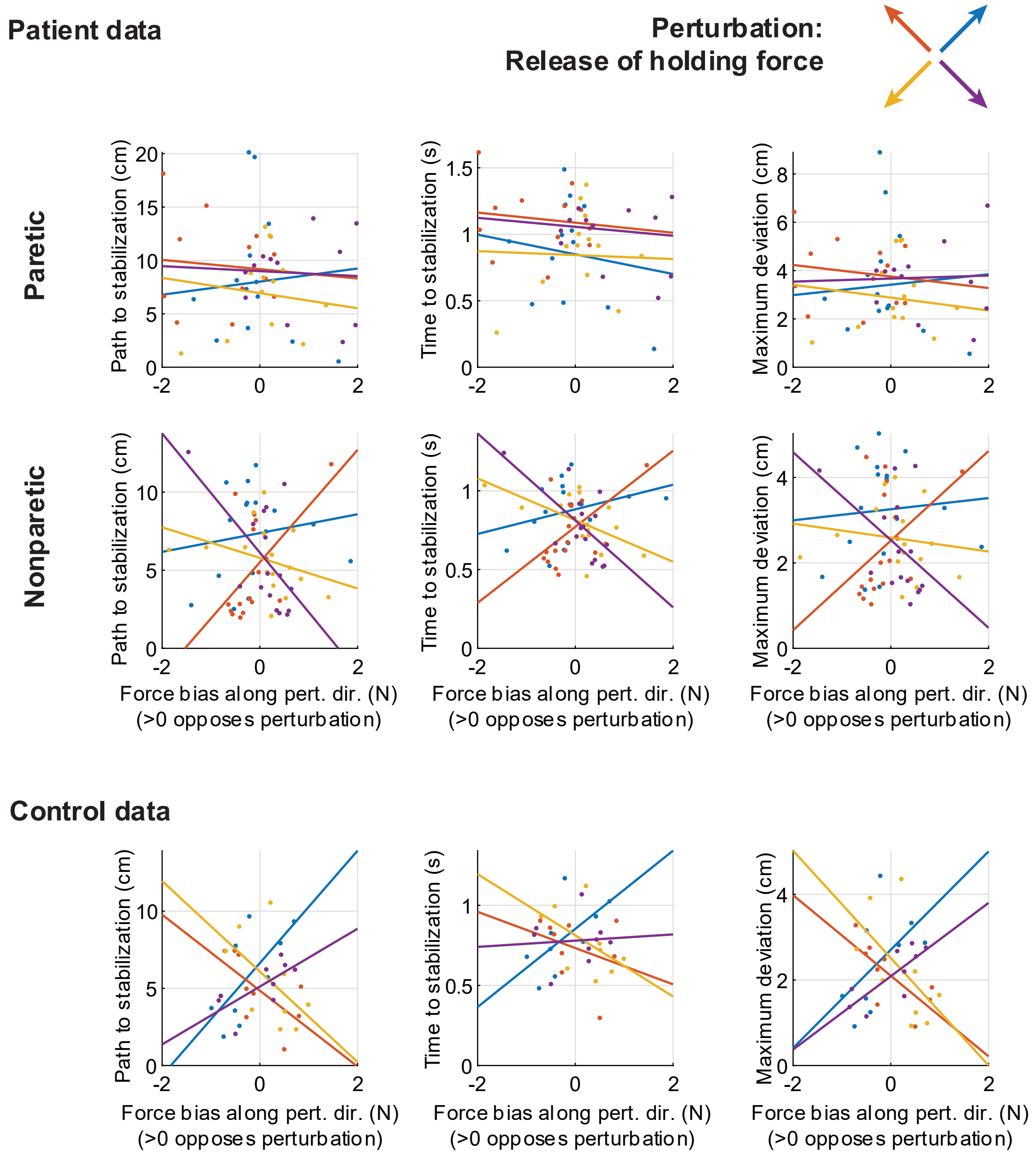


***Figure S7-3:*** *Across-patient comparisons between average postural biases (projected along the direction of the release perturbation) and different metrics of performance against the release perturbation: path to stabilization (left column); time to stabilization (middle column); and maximum deviation (right column). Different colors indicate data related to the four different release perturbation directions. Lines indicate linear fits. Positive values in the x-axis indicate biases that would oppose the perturbation. Note the limitations of these inter-individual analyses (which also hold for Figures S5-3 and S6-3): First, averaging these effects for each individual would average out opposing contributions of resting biases; here, because the perturbations come in exactly opposing pairs, this would average to zero, which is why we show these relationships for each perturbation separately instead (which may result in higher measurement noise). Second, the power of correlation analyses may be diluted by inter-individual differences in other factors, such as overall stiffness. Focusing on within-individual differences addresses both these issues.*

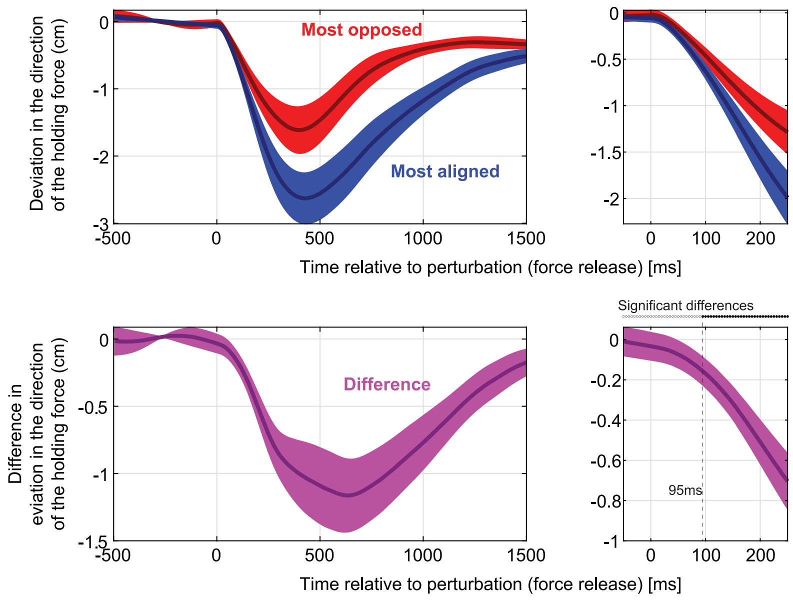


***Figure S7-4: Top:*** *Time course of responses to the static release perturbation for the cases where the resting bias was the most aligned (blue) vs. the most opposed (right) to the perturbation. The right panel zooms in the first 250ms following the release.* ***Bottom:*** *difference between the most-opposed vs. most-aligned data shown on top. On the right panel, statistically significant differences emerge 95ms after the release.*

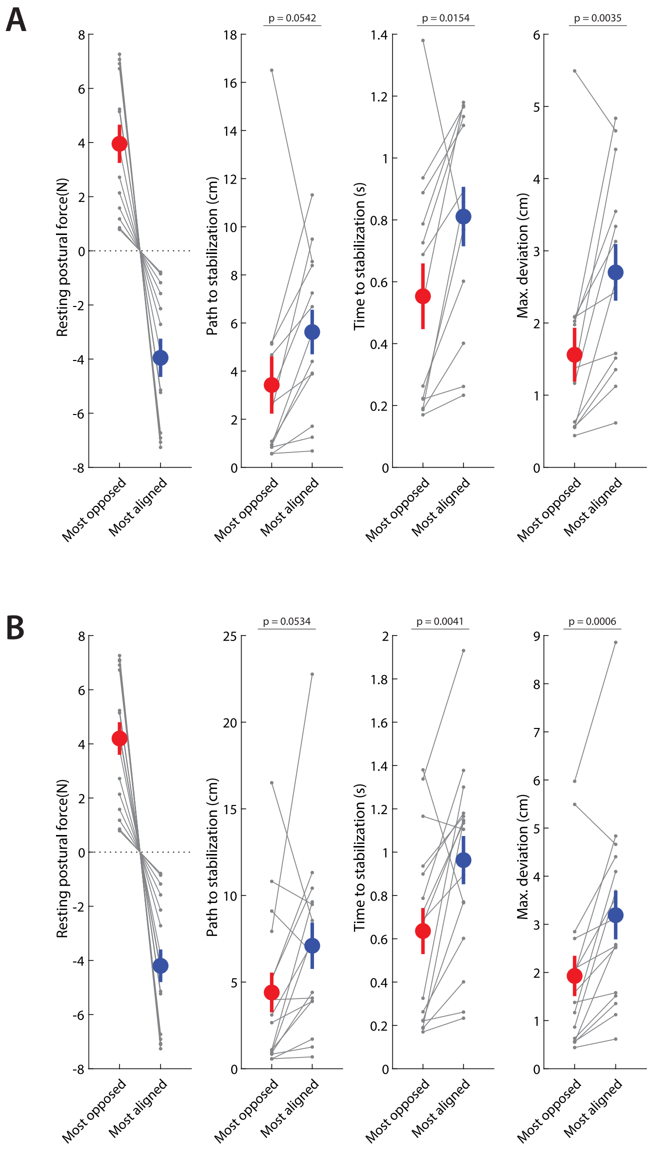


**Figure S7-5:** *Repeating the analyses in Figure 7F to ensure no systematic effects of missing data.* In some holding perturbation trials, patients took a long time to reach the stabilization criterion described in the previous section; mistakenly, our setup limited its recording time to only the first 2s after force release. The exact time to stabilization thus could not be measured for these particular trials, so they had to be excluded from analysis. Though only 13.2±3.3% (mean±SEM) of paretic stabilization trials were thus excluded in the patient population (1.4±0.4% in their non-paretic side, 0.4±0.4% [2 trials] in controls), there were three patients for which excluded trials were 25% or more of all paretic trials. To ensure there are no systematic effects of this issue, we repeated the analysis of Figure 7F (a) by excluding these three patients altogether (Shown in **A**) or (b) by assigning a value of 2.0 seconds to the affected trials. In both cases, we found results similar to our main analysis (Shown in **B**). Both analyses yielded results similar to our main analysis.

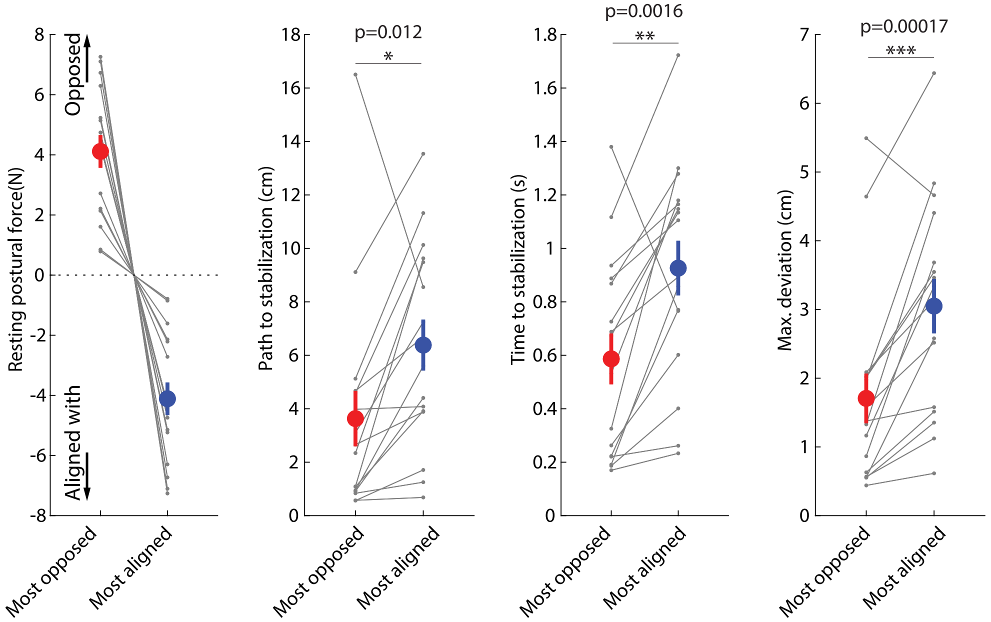


***Figure S7-E:*** *Repetition of the analysis in Figure 7F but with resting biases calculated without trial rejection, showing similar results (performance against the static release perturbation is better when the resting biases are directed against the perturbation, and worse when the resting biases are aligned with the perturbation, showing interaction between resting biases and active holding control).*
